# Bayesian inference of reassortment networks reveals fitness benefits of reassortment in human influenza viruses

**DOI:** 10.1101/726042

**Authors:** Nicola F. Müller, Ugnė Stolz, Gytis Dudas, Tanja Stadler, Timothy G. Vaughan

## Abstract

Reassortment is an important source of genetic diversity in segmented viruses and is the main source of novel pathogenic influenza viruses. Despite this, studying the reassortment process has been constrained by the lack of a coherent, model-based inference framework. We here introduce a novel coalescent based model that allows us to explicitly model the joint coalescent and reassortment process. In order to perform inference under this model, we present an efficient Markov chain Monte Carlo algorithm to sample rooted networks and the embedding of phylogenetic trees within networks. Together, these provide the means to jointly infer coalescent and reassortment rates with the reassortment network and the embedding of segments in that network from full genome sequence data. Studying reassortment patterns of different human influenza datasets, we find large differences in reassortment rates across different human influenza viruses. Additionally, we find that reassortment events predominantly occur on selectively fitter parts of reassortment networks showing that on a population level, reassortment positively contributes to the fitness of human influenza viruses.

## Introduction

Through rapid evolution, human influenza viruses are able to evade host immunity in populations around the globe. In addition to mutation, reassortment of the different segments of influenza viruses provides an important source of viral diversity (Steel and Lowen, 2014). If a cell is infected by more than one virus, progenitor viruses can carry segments from more than one parent (McDonald et al., 2016). with the exception of accidental release of antigenically lagged human influenza viruses (Nakajima, Desselberger, and Palese, 1978), reassortment remains the sole documented mechanism for generating pandemic influenza strains (e.g (Smith, Bahl, et al., 2009; Smith, Vijaykrishna, et al., 2009; Guan et al., 2010)).

To characterize such events, tanglegrams, comparison between tree heights (Westgeest et al., 2014; Dudas, Bedford, et al., 2014), or ancestral state reconstructions (Lu, Lycett, and Brown, 2014) are typically deployed. These approaches identify discordance between different segment tree topologies or differences in pairwise distances between isolates across segment trees. Tanglegrams in particular require a substantial amount of subjectivity and have been described as potentially missleading (De Vienne, 2018).

While the reassortment process has been intensively studied (e.g (Nelson et al., 2008; Westgeest et al., 2014; Dudas, Bedford, et al., 2014; Lu, Lycett, and Brown, 2014)), there is currently no explicit model based inference approach available. We address this void by introducing a coalescent-based model of the reassortment of viral lineages. In this phylogenetic network model, ancestral lineages carry genome segments, of which only a subset may be ancestral to sampled viral genomes. As in a normal coalescent process, network lineages coalesce (merge) with each other backwards in time at a rate inversely proportional to the effective population size. We model reassortment (splitting) events as a result of a constant-rate Poisson process on network lineages. At such a splitting event, the ancestry of segments on the original lineage diverges, with a random subset following each new lineage. We thus explicitly model reassortment networks and the embedding of segment trees within these, allowing us to infer these entities from available sequence data.

In order to perform inference under such a model, the reassortment network and the embedding of each segment tree within that network must be jointly inferred. This is similar to the well-known and challenging problem of inferring ancestral recombination graphs (ARGs), with the difference that segments in our model have fixed boundaries, but no defined ordering. While many approaches to inferring ARGs exist, some are restricted to tree-based networks (Didelot et al., 2010; Vaughan, Welch, et al., 2017), meaning that the networks consist of a base tree where recombination edges always attach to edges on the base tree. Other approaches (e.g (M. D. Rasmussen et al., 2014)) rely on approximations (McVean and Cardin, 2005) and are not applicable to the reassortment model due to the aforementioned lack of segment ordering. Completely general inference methods exist (Erik W. Bloomquist and Marc A. Suchard, 2010), but these are again not directly applicable to modelling reassortment and furthermore tend to be highly computationally demanding.

Here we introduce a novel Markov Chain Monte Carlo (MCMC) approach to jointly sample networks and the embedding of segment trees within those networks, without any approximations involved. This approach allows us to perform joint inference of the reassortment network, the phylogenetic trees of each segment, the reassortment and coalescent rates with the evolutionary models and its parameters.

We first show that this approach is able to retrieve reassortment rates, effective population sizes and reassortment events from simulated data. Secondly, we discuss how a lack of genetic information influences the inference of these parameters. Thirdly, we show how using the coalescent with reassortment can influence the inference of effective population sizes, as well as evolutionary rates. We then apply this approach to study reassortment rates patterns across different influenza subtypes. Finally, we study how reassortment rates differ on fit and unfit edges of these reassortment networks.

### Inference of effective population sizes and reassortment rates are reliably inferred from genetic sequence data

In order to test our ability to infer effective population sizes and reassortment rates from genetic sequences, we performed a well calibrated simulation study. To do so, we first sampled random effective population sizes from a log normal distribution (mean 5 and standard deviation 0.5) and reassortment rates from another log normal distribution (mean 0.2 and standard deviation 0.5). We then sampled the sampling times of 100 taxa, each with 4 segments, from a uniform distribution between 0 and 20. We next simulated reassortment networks alongside the embedding of the segment trees using these parameters. For each segment tree, we next simulated genetic sequences by using the JC69 substitution model (Jukes and Cantor, 1969) with an evolutionary rate of 5 × 10^-3^ per site and year. Each segment thereby consisted of 1000 independently evolving nucleotides. In order to study the effect of reducing the amount of genetic information, we additionally considered the scenario where all segments had an evolutionary rate of 5 × 10^-4^ per site and year. Using our MCMC approach we then inferred the reassortment network, segment tree embedding, effective population sizes and reassortment rates from these genetic sequences.

The results shown in figure 1A,B, indicate that we are able to correctly retrieve effective population sizes and reassortment rates from simulated genetic sequences. Effective population sizes are estimated more precisely than reassortment rates, which is expected considering that there are typically many more coalescent events in a network than reassortment events. Lower evolutionary rates do not greatly decrease our ability to infer effective population sizes and reassortment rates (see figure S1).

**Figure 1:**
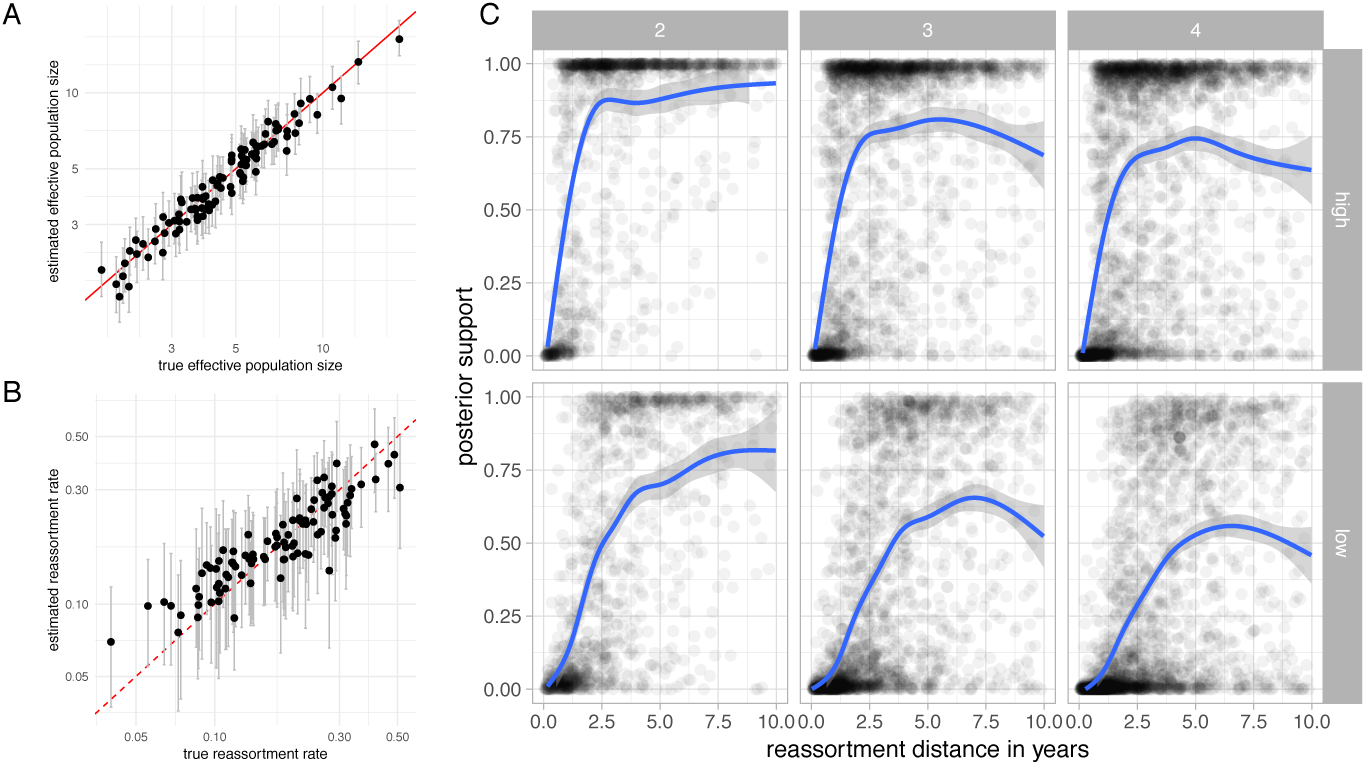
Estimates of effective population sizes and reassortment rates from simulated genetic sequences. **A** Estimated effective population sizes and 95% confidence intervals (y-axis) vs. simulated effective population sizes on the x-axis. **B** Estimated reassortment rates and 95% confidence (y-axis) vs. simulated reassortment rates on the x-axis. **C** Posterior support for true reassortment events (y-axis) given the reassortment distance (x-axis). Inference of reassortment networks from sequences simulated with a evolutionary rate of 5 × 10^-3^ mutations per site and year (top row) and 5 × 10^-4^ mutations per site and year (bottom row). From left to right, the reassortment events are for networks with 2,3 and 4 segments.

To test how well true reassortment events are recovered, we computed the probability of observing exactly the same reassortment events as present in the true (simulated) network. We considered to reassortment events to be the same if the sub-tree of each segment below that node is the same and if the relative direction of each segment at the reassortment event is exactly the same (see Methods, Network Summary). This constitutes a stringent definition of two reassortment events being the same.

As shown in figure 1C, reassortment events are well supported, particularly with increasing reassortment distance. The reassortment distance denotes how much independent evolution happened on the two parent viruses of the reassortment event (see Methods, Reassortment Distance).

This is particularly true when we only look at reassortment events between pairs of segments and drops when we look at 3 or 4 segments. This decrease is driven by our definition that two reassortment events are only the same if all segments reassort in the same relative direction at the same time with exactly the same clade below the segment trees; a requirement that becomes harder to satisfy as the number of segments increases. As expected for methods that correctly take into account uncertainty, the posterior support decreases when lower evolutionary rates are used to simulate the sequences of the segments.

### Joint inference from full genomes increases precision in dating nodes

We compared the internal node ages inferred using the coalescent with reassortment to ages inferred under the assumption that all segments evolved independently under the standard coalescent model. To do this, we first compiled datasets of several human influenza A subtypes, as well as influenza B (details in the Materials and Methods). From each of these we generated three datasets consisting of a random sample of sequences. We then analysed each of these sub-sampled datasets once using the coalescent with reassortment and once using a normal coalescent prior with shared effective population size across all segments, but assuming that each segment evolved independently. We first computed the 95% highest posterior density (HPD) interval of node heights for each clade that was supported by both approaches with a posterior probability of more than 0.5. We then normalized the difference between the lower and upper bound of the 95% HPD interval, by the median node height estimate to get the relative with of the HPD interval for each clade. As shown in figure 2A, using the coalescent with reassortment reduces the uncertainty of node height estimates of segment tree nodes by 34% for p2009 like H1N1)up to 50% for influenza B.

**Figure 2:**
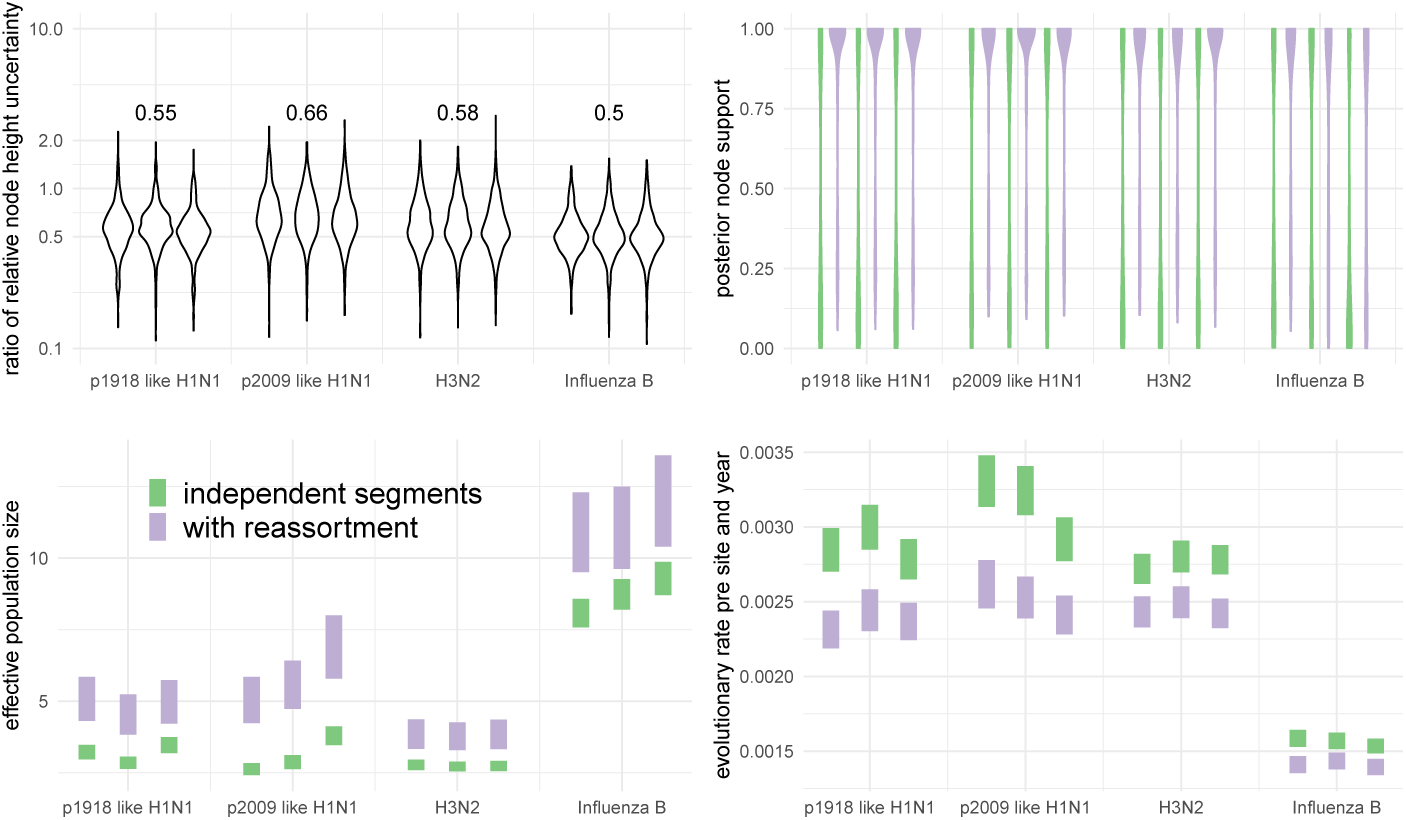
Comparison of estimates between the coalescent with reassortment and assuming that each segment codes for an independent realization of the same coalescent process. **A** Comparison of the relative width of the 95% HPD interval of segment tree node heights. The vertical axis shows the distribution of ratios of the relative width of the 95% HPD intervals of the coalescent with reassortment over the coalescent assuming independent segment evolution. The values show the median reduction in node height uncertainty when using the coalescent with reassortment over the coalescent with independent segments. **B** Comparison between the distribution of posterior clade support of segment trees found the maximum clade credibility segment trees. **C** Comparison between the inferred effective population sizes. When assuming each segment is an independent realization of the same coalescent process, the effective population sizes are inferred to be much smaller and much more certain. **D** Comparison between the inferred clock rates. The coalescent with reassortment infers lower clock rates.

Next we computed the distribution of clade supports for clades represented in the MCC trees inferred using the two approaches. As shown in figure 2B, segment tree clades are much better resolved when using the coalescent with reassortment for all datasets.

We then compared the effective population sizes and evolutionary inferred using the two approaches. The coalescent with reassortment infers higher effective population sizes for all datasets (see figure 2C). This also influences the inferred clock rates, since lower effective population sizes put stronger weight on shorted branches and therefore larger clock rates (see figure 2C). We explain this discrepancy as follows. Coalescent events closer to the tips are more likely between lineages that are for example geographically more closely related and can be assumed to occur rapidly and provide information about low effective population size values. Coalescent events deeper in the tree on the other hand are more representative of those between geographically more separated lineages. These events therefore provide information about larger effective population sizes. Coalescent events across different segments that occur close to the tips are less likely to have encountered reassortment events. In the coalescent with reassortment, they are therefore interpreted as one event, whereas in the coalescent with independent segments, they are interpreted as eight. Coalescent events deeper in the tree are more likely between lineages that encountered reassortment events and are therefore more likely to provide independent information about the population process. The coalescent with independent segments assumes that all coalescent events provide the same amount of information about the population process and will consequently favour information about the population process closer to the tips. This leads to differences in the estimated effective population sizes which then leads to differences in the estimated clock rates.

We also compared the performance of the two approaches by inferring tip dates (Dudas and Bedford, 2019). The tips (leaf nodes) are the only nodes in the trees or network for which we can actually presume to know the true age, which is set by the sample collection time. To compare the two approaches, we compiled 1000 smaller influenza A/H3N2 datasets each composed of 20 genomes. Of those datasets, 500 were randomly sampled from an interval of 2 years between 1995 and 2019. The remaining 500 datasets were assembled using a random sampling interval of 10 years between 1995 and 2019. From each of these datasets we randomly selected a single genome and inferred its sampling time using both approaches, conditional on the sampling times of the remaining genomes.

The 95% HPD of the sample time posteriors under the coalescent with reassortment contains the true sampling time interval in 91% of cases for the 2 year sampling interval and in 89% for the 10 year sampling interval (see figure S2). On the other hand, the 95% HPD of the sample time posteriors generated by the independent segment coalescent model contains the true sampling time in only 68% (2 year sampling interval) and 77% (10 year sampling interval) of cases.

### Contrasting reassortment rates across different human influenza viruses

We compared the reassortment rates of different influenza types. To do so, we used the same datasets as described above, as well as an influenza A/H2N2 dataset sampled between 1957 and 1970. We then jointly inferred the reassortment network, the embedding of segment trees, evolutionary rates, effective population sizes and reassortment rates of these viruses. We find that the estimated reassortment rates vary greatly between different influenza viruses.

Influenza A/H3N2 shows the highest rates of reassortment, while pandemic 1918 like H1N1 and influenza B show the lowest inferred rates of reassortment (see figure 3). H2N2 and 2009 pandemic like H1N1 show intermediate rates of reassortment, although the uncertainty on those estimates is quite large.

**Figure 3:**
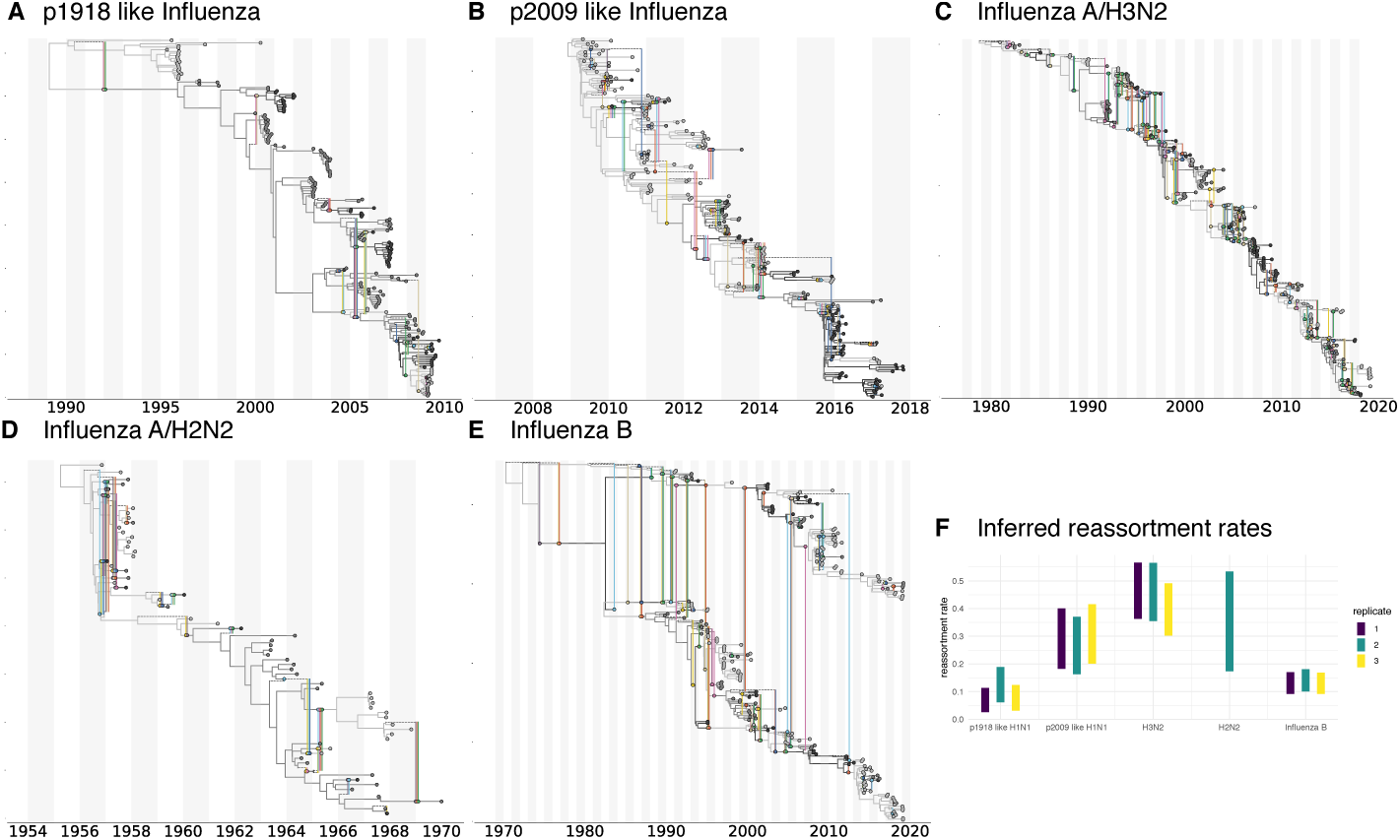
Estimates of maximum clade credibility networks and reassortment rates of different human influenza viruses. Maximum clade credibility (mcc) networks of p1918 Influenza A/H1N1 **A**, p2009 influenza A/H1N1 **B**, influenza A/H2N2 **C**, influenza A/H3N2 **D** and influenza B **E**. These mcc networks are show for one of the random subsets. Rhe mcc networks of all random subsets are shown in figures S3-S6. **F** Here we show the inferred reassortment rates (y-axis) for different influenza viruses on the x-axis. The reassortment rates are per lineage and year.

Differences between p1918-like H1N1, H2N2 and H3N2 are particularly interesting since these strains share many common segments. All segments with the exception of HA, NA, and PB1 of influenza A/H3N2 originate from the p1918-like H1N1 strain (Scholtissek et al., 1978) and H2N2 and H3N2 only differ in HA and PB1. Pandemic 2009-like human H1N1 which became seasonal in the years after the 2009 pandemic on the other hand has one segment (PB1) that originates from human H3N2 and three segments (HA, NP, and NS) derived from classic swine viruses which are descended from a p1918-like strain (Smith, Vijaykrishna, et al., 2009). It shows similar reassortment rates to H3N2, but highly elevated levels compared to the p1918-like H1N1 strain.

Such variations in reassortment rates may be driven by a number of factors. Differences in co-infection rate (which may be linked to the effective population size) lead to different probabilities of viruses being in the same host at the same time and therefore to difference in the rate at which reassortants appear. In particular, the higher incidence of Influenza A/H3N2 and the correspondingly likely higher number of co-infection events compared to other influenza A viruses or influenza B viruses may contribute to the higher observed reassortment rate in that case. Additionally, potentially different survival probabilities of reassortants could affect the observed reassortment rates.

### Reassortment events occur on fitter parts of reassortment networks

Next, we test if there is a fitness effect associated with reassortment events. To do so, we classify every network edge from the posterior distribution of inferred networks as either “fit” or “unfit”. We define a fit edge to be any edge having descendants which still persist at least 2 years into the future, while every other edge in the network is defined to be unfit. If reassortment events are beneficial, lineages that are the result of reassortment events should have a higher survival probability and are therefore more likely to persist further into the future.

To test if this is the case, we calculated the number of reassortment events on fit edges and on non-fit edges for all networks in the posterior distribution of the MCMC. We then divided this number by the total length of fit respectively not fit edges. As shown in figure 4A, reassortment events occur at a higher rate on fit edges of the H3N2 and influenza B networks than they do on non-fit edges. This suggests that reassortment is beneficial to the fitness of influenza A/H3N2 and influenza B viruses.

**Figure 4:**
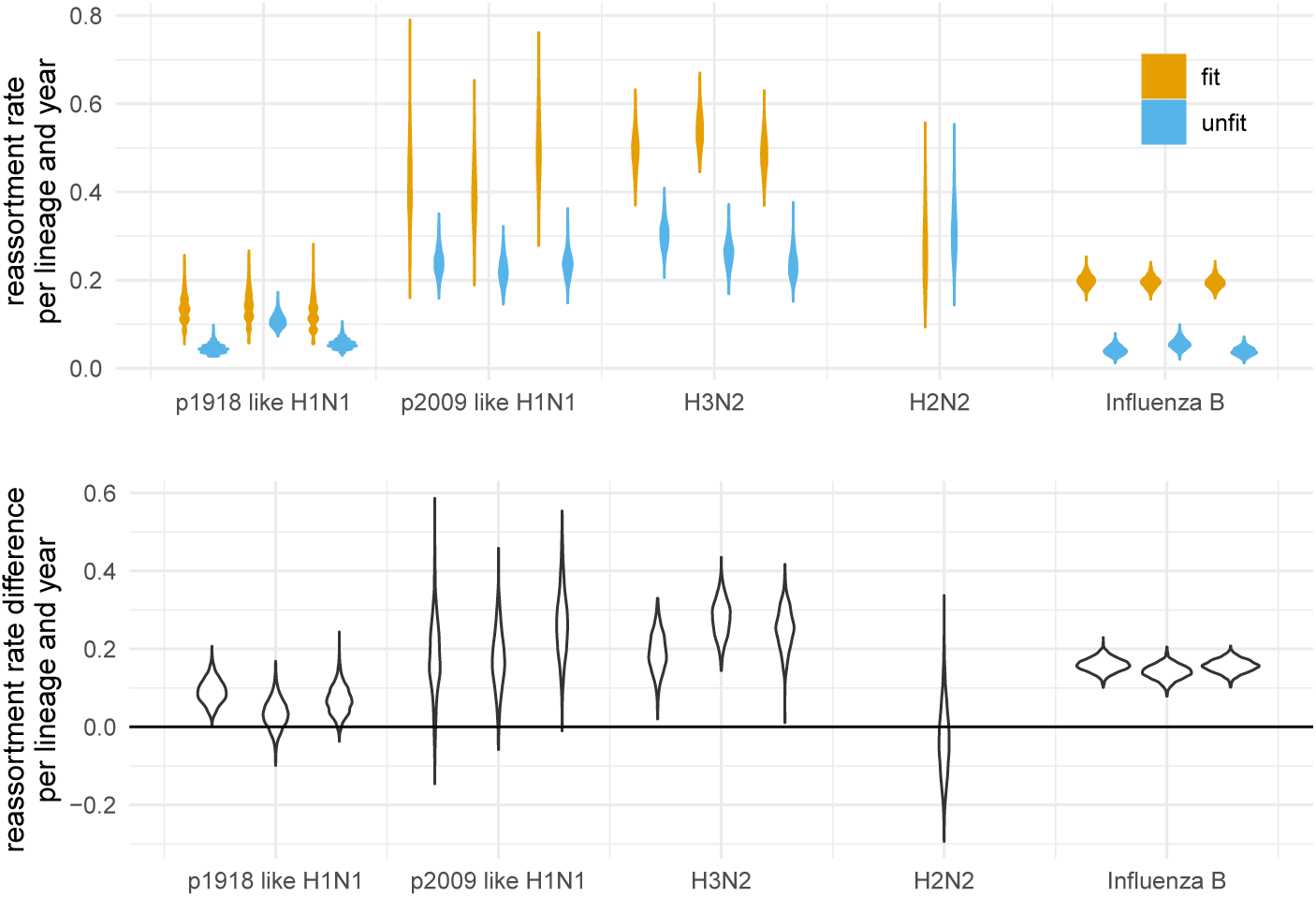
Estimates of reassortment rates on fit and unfit edges. **A** Here we show the number of reassortment events on fit and unfit edges of the networks divided by the total length of fit and unfit edges. Fit edges are defined as having sampled descendant at least 2 years into the future. Every other edge is considered unfit. These rates are shown for different human influenza viruses on the x-axis. The violin plots denote the distribution of theses ratios over the posterior distribution of networks. **B** Here we show the difference between fit and unfit reassortment rates. Values above 0 indicate that reassortment events are more likely to occur on fitter, while values below 0 indicate that reassortment events are more likely to occur less fit edges.

For the other human influenza viruses, fitness benefits of reassortment are less pronounced. For p09-like H1N1 and H2N2, the sampling time windows (both) or number of samples (H2N2) was however rather small and the results are likely driven by a lack of data. Non p09-like H1N1 on the other hand had relatively few reassortment events overall driven by a low reassortment rate and the results are likely driven by a lack of reassortment events. These patterns largely hold if the definition of what is a fit edges is changed to having descendants at least 4 or 6 years into the future (see figure S7 & S8). It however decreases for p09 like H1N1 for which the overall sampling interval is only 10 years.

Since H3N2 has been densely sampled over long time intervals, we analysed two more influenza A/H3N2 datasets, one sampled between 1980 and 2005 and one sampled from 2005 until today. For both these datasets, we find higher rates of reassortment on fit edges (see figure S9). We next tested if for datasets sampled over short times (2 years), we would estimate reassortment rates consistent with the estimated rates on unfit edges. To do so, we compiled 9 dataset, each with 100 to 200 sequences sampled from 2 seasons between 2000 and 2018. Averaged over all 9 datasets, we find the short-term reassortment rate to be approximately 0.2 reassortment events per lineage per year, which is consistent with the reassortment rate estimates for unfit edges (see figure S10).

Finally, we sought to rule out the possibility that these patterns are simply a property of our reassortment model. To do this, we simulated networks under the coalescent with reassortment with the reassortment rates and effective population sizes fixed to the mean values estimated from the empirical data, and the network leaf times fixed to those from the same data. We then recomputed the same fit/unfit reassortment rate statistics from these simulated networks (see figure S11) and found that the patterns we observed in the empirical data observed patterns completely disappeared. This strongly suggests that the elevated rate of reassortment on fit lineages is not due to the particulars of our model, but is instead a real effect.

## Conclusion

We here present a novel Bayesian approach to jointly infer the reassortment network, the embedding of segment trees and the corresponding evolutionary parameters. We show that this approach is able to retrieve reassortment rates, effective population size and reassortment events from simulated data.

We have used this facility to show that there are larger differences in the rates of reassortment across different Influenza viruses, and that reassortment events occur predominantly in fitter parts of the corresponding reassortment networks. We propose that this is due to selection favouring lineages that have reassorted. Although we have deployed a relatively simple way of defining which edges of a network are fit and which are not, future approaches could more directly incorporate fitness models into these network type approaches (Łuksza and Lässig, 2014; Neher, Russell, and Shraiman, 2014).

Even if one is not directly interested in reassortment patterns, our approach allows phylogenetic and phylodynamic inferences to exploit full genome sequences for reassorting viruses. This helps to avoid bias and increase precision compared to, for instance, assuming segments evolve completely independently. However, a lot of development remains to be done in the direction of incorporating skyline models for population size dynamics (Drummond et al., 2005; Minin, Erik W Bloomquist, and Marc A Suchard, 2008) together with extending the model to account for population structure (Vaughan, Kühnert, et al., 2014; Müller, D. A. Rasmussen, and Stadler, 2017).

In summary, this approach allows us to perform network inference by directly accounting for a special kind of recombination process, i.e. reassortment. In the future, we will pursue the development of related approaches to account for a variety of other recombination processes.

## Methods and Materials

### The coalescent with reassortment

Here we introduce a model to describe a coalescent process with reassortment. To do so, we define *t* to be the time (increasing into the past) before the most recent sample, and *L*_*t*_ as the set of network lineages extant at time *t* (see Figure 5). Each extant network lineage *l ∈ L*_*t*_ carries the full set of genome segments, *S*. In general however, only a subset *C*(*l*) ⊆ *S* of these are directly ancestral to sampled viruses. We refer to this subset as the *carrying load*. We further define the total number of segments |*S*| and the number of ancestral segments |𝒞(*l*)|.

**Figure 5:**
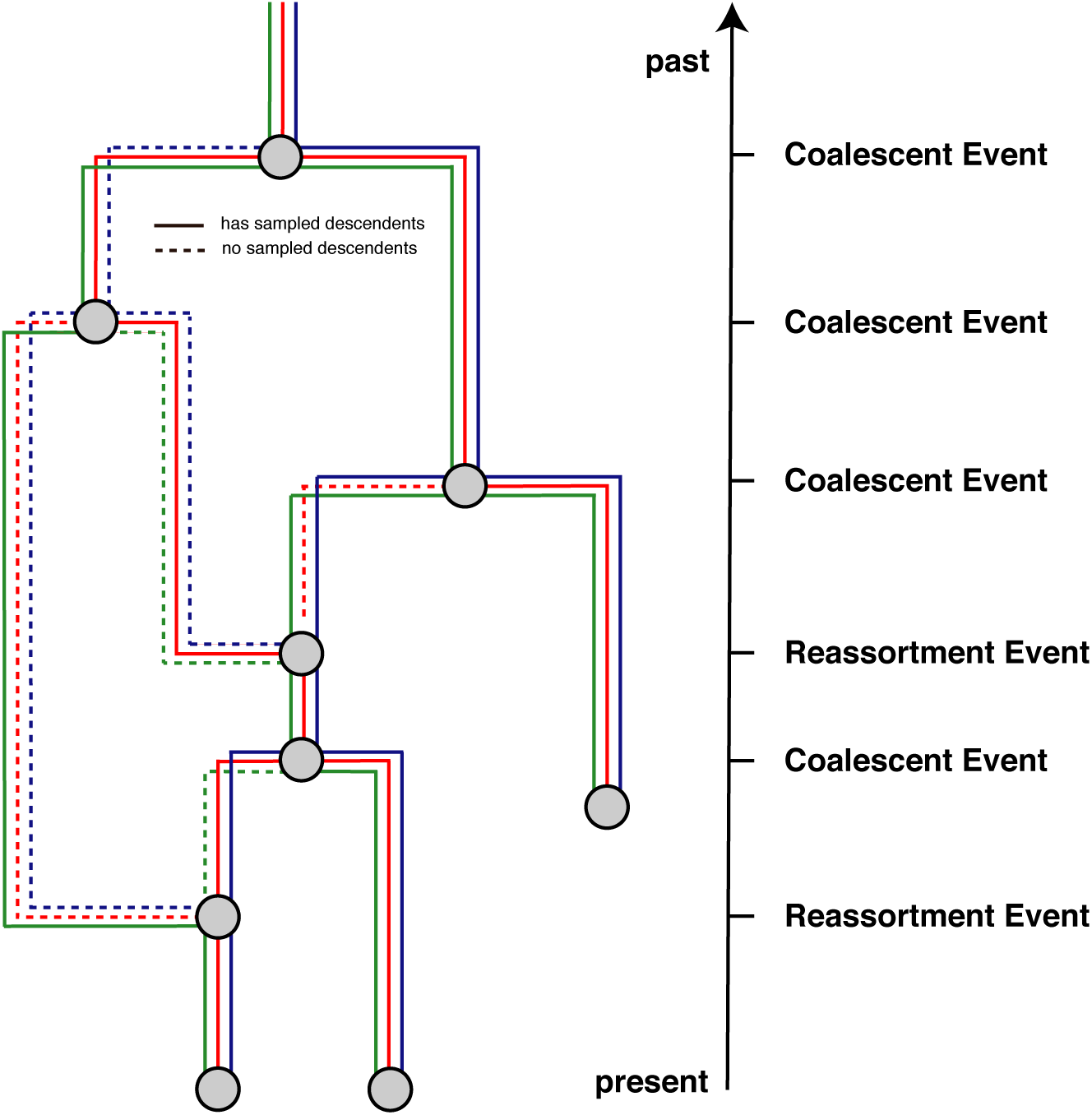
Example reassortment network. Here we give an example of a reassortment network where we track 3 different segments differentiated by the different colors through the network. Dashed lines denote segment lineages that do not have sampled descendants. As done in coalescent approaches, we track the network from the present backwards in time to the past.

The coalescent with reassortment is a continuous time Markov process that proceeds backward in time. It involves three possible events: *sampling, coalescent* and *reassortment* events. As is usually case for coalescent approaches, we condition on sampling events. These happen at predefined times and simply the number of active network lineages by 1. Coalescent events occur between two network lineages *l* and *l’* at a rate that is inversely proportional to the effective population size *N*_*e*_ and reduce the number of active network lineages by 1. The smaller the effective population size, the more likely two lineages are to share a common ancestor, i.e. the more likely they are to coalesce. Upon a coalescent event, the segments that the parent lineage *p* of lineages *l* and *l’* carries is the union of the segments that is carried by *i* and *j*, i.e.:

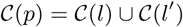

This coalescent events in the network only corresponds to a coalescent event in a segment tree when the corresponding segment is present in both *C*(*l*) and *C*(*l’*).

Reassortment events happen at a rate *ρ* per lineage per unit time. A reassortment event on lineage *l* will increase the number of network lineages by 1. The segments carried by lineage *l* are randomly assigned to the two parent lineages *p*_1_ and *p*_2_. This means that the probability of the ancestry of a given segment to follow *p*_1_, for example, is 0.5.

As we are not interested in the history of segments that are not ancestral to our sample, we explicitly integrate over this ancestry in our model. As with standard coalescent with recombination models, this is done by omitting non-ancestral events from the process and modifying the reassortment rate to exactly account for this omission. In our model, the events which are omitted are “reassortment” events on *l* in which the ancestry of every ancestral lineage in *C*(*l*) is assigned to the same parent. (Thus no true reassortment occurs.) Since each segment chooses its parent edge uniformly at random, the probability of either *p*_1_ or *p*_2_ being chosen as ancestral to all segments is

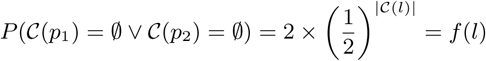

The effective rate of “observable” reassortments on lineage *l* is then simply *ρ*(1 - *f* (*l*)).

### Calculating the posterior probability

In order to perform joint Bayesian inference of reassortment networks together with the parameters of the associated models, we use a Markov chain Monte Carlo (MCMC) algorithm to characterize the joint posterior density

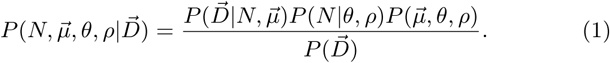

Here *N* represents the full reassortment network (including the embedding of the segment trees), the elements of the vectors 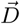 and 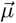 represent the segment-specific multiple sequence alignments and their associated molecular substitution models and parameters. The parameters *θ* and *ρ* are the effective population size and per-lineage reassortment rate.

The terms on the right-hand side of Eq. (1) include the network like-lihood 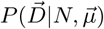, the network prior *P (N\ρ, θ)* and the joint parameter prior 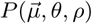. Each of these terms is discussed below. (The denominator 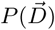 is the marginal likelihood of the model and does not concern us here.)

### The network likelihood

The usual conditional independence of sites assumption made in phylogenetic analyses allows us to factorize the network likelihood in terms of the individual segment tree likelihoods:

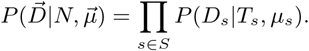

These tree likelihoods can be computed using the standard pruning algorithm (Felsenstein, 1981).

### The network prior

The term *P* (*N* |*θ, ρ*) denotes the probability of the network and the embedding of segment trees under the coalescent with reassortment model, with effective population size *θ* and per-lineage reassortment rate *ρ*. It plays the role of the tree prior in standard phylodynamic analyses.

We can calculate *P* (*N* |*θ, ρ*) by expressing it as the product of exponential waiting times between reassortment, coalescent and sampling events, i.e.:

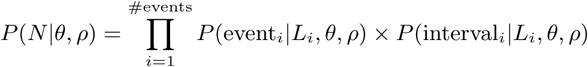

where we define *t*_*i*_ to be the time of the *i*^th^ event, and *L*_*i*_ to be the set of lineages extant immediately prior to this event. (That is, *L*_*i*_ = *L*_*t*_ for *t ∈* [*t*_*i-*1_, *t*_*i*_).)

### Event contribution

The event contribution of the *i*^*th*^ event in the network is different depending on if the *i*^*th*^ event is a coalescent or reassortment event. If the *i*^*th*^ event is a coalescent event between lineage *l*_1_ and *l*_2_, the event contribution is the probability density of this particular pair of lineages coalescing at time *t*_*i*_. For a constant-sized coalescent model, this is

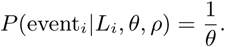

On the other hand, if the *i*^*th*^ event is a reassortment event on lineage *l*, the event contribution is the probability density of an (observable) reassortment event to occur on that lineage, i.e:

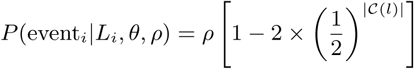

As we condition on sampling events, their event contribution is always simply 1.

### Interval contribution

The interval contribution *P* (interval_*i*_|*L*_*i*_, *θ, ρ*) is the probability of not observing any event in a given time interval. Three different types of events can happen in the coalescent with reassortment: sampling, coalescent and reassortment events. Since we condition on the times of the sampling events, only coalescent and reassortment events are produced by the CTMC. Given the total rate Λ_*i*_ (probability per unity time) with which these occur in the interval immediately prior to event *i*, the interval contribution can be written as

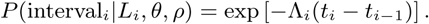

The total rate is the sum of the coalescence rate 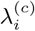 and the reassortment rate 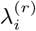. The coalescence rate depends on the number of lineages extant at a particular time and the effective population size in the usual way.

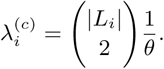

The rate of observable reassortment events is

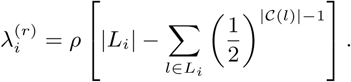

Note that this is generally less than the total rate of reassortment events in this interval, which would be simply *ρ*|*L*_*i*_|, as this rate excludes reassortment events that produce lineages carrying no ancestral segments.

### The parameter priors

The term 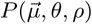 denotes joint prior distribution of all model parameters. We factorize this, writing it as the product of the individual parameter priors 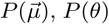 and *P* (*ρ*). This asserts that our prior information on any one of these model parameters is independent of the prior information we have for the others.

## An MCMC algorithm for reassortment networks

In order to perform MCMC sampling of network and the embedding of segment trees within these networks, we introduce several MCMC operators. These operators often have analogues in operators used to explore different phylogenetic trees. We here only briefly discuss what these operators are continually doing and provide more details in the supplement:

1. Add/Remove operators which add and remove reassortment events extends the SPR move for networks (Bordewich, Linz, and Semple, 2017) to jointly operate on segment trees as well.
2. Segment diversion operators which change the path segments take at reassortment events.
3. Exchange operators which change the attachment of edges in the network while keeping the network length constant.
4. Sub-network Slide operators which change the height of nodes in the network while allowing to change the topology
5. Scale operators which scale the heights of individual nodes or the whole network without changing the network topology.
6. Re-simulating above the segment tree roots operator, to efficiently sample parts of the network that are not informed by any genetic data.
7. Empty segment operator to augment the network with edges that do not carry any segments for the duration of a move, to allow larger jumps in network space.

We validate the the implementation of the coalescent with reassortment network prior as well as all operators in the supplement.

## Summarizing reassortment networks

To summarize over a distribution of networks, we use a similar strategy to the maximum clade credibility strategy used to summarize over distributions of trees. To do so, we first compute all unique coalescent and reassortment nodes was encountered during the MCMC. To do so, we have to define when two coalescent or reassortment nodes are the same. We define two coalescent nodes to be the same if a) the parent edges of those nodes carry the same segments and b) if the sub-tree below each segment includes exactly the same clades between the two coalescent nodes. We define two reassortment nodes to be the same if a) both parent edges carry the same segments in the same relative orientation and b) if the sub-tree below each segment includes exactly the same clades between the two reassortment events. This however also means that the more segments we include in the summary, the more likely two nodes will be considered different nodes.

While the number of coalescent and reassortment nodes in the network changes over the course of the MCMC, the number of coalescent nodes on the segment trees is constant. In order to avoid dimensionality issues when summarizing, we first compute the frequency of observing each coalescent node over the course of the MCMC. We then weight this frequency by the number of coalescent events on segment trees this coalescent node corresponds to. We next choose the network that maximizes those weighted clade credibilities as the maximum clade credibility (or MCC) network.

In order to compute the posterior support of each reassortment event in the MCC network, we next compute the frequency of observing each reassortment event in the MCC network during the MCMC.

Since we require the network to be rooted, we track segments event after the root of a segment tree was reached. These patterns are however not supported by any genetic information and follow the prior distribution only. For the summary of networks, we therefore remove segments from edges if the root of a segment tree has been reached. Additionally, we remove reassortment loops, i.e. events that start on one edge and then directly reattach to the same edge. Since the support for individual events can greatly depend on how many segments are analysed, we also implemented the option to only summarize over a subset of the segments, while ignoring others.

## Reassortment distance

For any reassortment event where segments *a* and *b* take different paths, we compute the reassortment distance of segment *a* onto segment *b* as follows: First, we follow segment *a* until it reaches a network edge that carries segment *b*. We then compute the common ancestor height between segment *b* at the reassortment event and segment *b* on that network edge. The reassortment distance of segment *a* onto segment *b* is then the different between this common ancestor height and the height of the reassortment event. This seeks to denote for how long segment *b* in the two parent viruses at the reassortment event evolved independently.

## Implementation

We implemented the MCMC framework for the Coalescent with Reassortment as a BEAST2 package called CoalRe. This package includes the classes to do simulation and inference under the coalescent with reassortment. The implementation is such that the tree likelihood calculations are separate from the the network framework, which allows to make use of the vast amount of different site and clock models implemented in BEAST2. Additionally, it can be used with other Bayesian approach such as Nested Sampling or coupled MCMC. Further, model comparison as well as integration over evolutionary models can be performed. The package can be downloaded by using the package manager in BEAUti. The source code for the software package can be found here: https://github.com/nicfel/CoalRe. A tutorial on how to set-up an analysis using the coalescent with reassortment is available at https://taming-the-beast.org/tutorials/Reassortment-Tutorial/ (Barido-Sottani et al., 2017). Networks are logged in the extended Newick format (Cardona, Rosselló, and Valiente, 2008) and can be visualized using for example icytree.org (Vaughan, 2017). Additionally, we provide python scripts to plot networks based on https://github.com/evogytis/baltic.

## Datasets and data availability

We compiled datasets from several influenza viruses using sequence data downloaded from fludb.org (pandemic and seasonal H1N1, H3N2 and influenza B). For the influenza A/H2N2 dataset, we ended up using the same sequences as in (Joseph et al., 2015). We downloaded these sequences from gisaid.org (acknowledgement table can be found here) For all datasets, but the influenza A/H2N2 dataset, we first sub-sampled all sequences to end up with at least 500 samples sampled evenly over time. We then aligned all segments using Muscle 3.8.31 (Edgar, 2004).

We then analysed every influenza virus under the coalescent with reassortment in BEAST 2.5.2 (Bouckaert et al., 2019) using coupled MCMC (Altekar et al., 2004; Mueller and Bouckaert, 2019). We assumed the sequences to have evolved under an HKY + Γ_4_ model (Hasegawa, Kishino, and Yano, 1985; Yang, 1993), allowing the first two codon position and the third having different rates (Shapiro, Rambaut, and Drummond, 2005). We then jointly estimated all evolutionary rates, the reassortment networks and embedding of segments trees, as well as the reassortment rates and effective population sizes. For the influenza A/H2N2 dataset, we additionally estimated the sampling times for all sequences for which only the year in which the sample was taken was known.

For virus types with sequences downloaded from fludb.org, the full XML files to run the datasets are available online. For the influenza A/H2N2 sequences that were obtained from gisaid.org, we remove the sequence characters from the XML files in order to comply with licence regulations of gisaid.org. Apart from the sequence characters, the XML files are complete. All other data, such as log files of BEAST2 runs, as well as scripts to analyse and plot results are available here https://github.com/nicfel/Reassortment-Material.

## Acknowledgement

We would like to thank Alexei J. Drummond, Simone Linz and Daniel Huson for useful discussions on how to summarize networks. NFM and TS are funded by the Swiss National Science foundation (SNF; grant number CR32I3 166258).

