## Supplement for "Bayesian inference of reassortment networks reveals fitness benefits of reassortment in human influenza viruses"

August 2, 2019

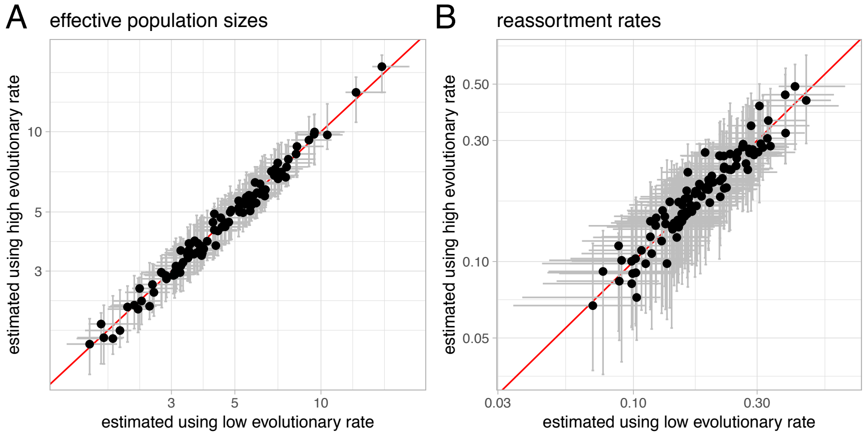


Figure S1: **Estimates of effective population sizes and reassortment rates from simulated genetic sequences. A** Estimated effective population sizes and 95% confidence intervals (y-axis) vs. simulated effective population sizes on the x-axis. The red diagonal depicts perfect equality. **B** Estimated reassortment rates and 95% confidence (y-axis) vs. simulated reassortment rates on the x-axis. We estimated these values from genetic sequences from 100 taxa from 4 segments that we simulated along segment trees of reassortment networks simulated using the parameters on the x-axis.

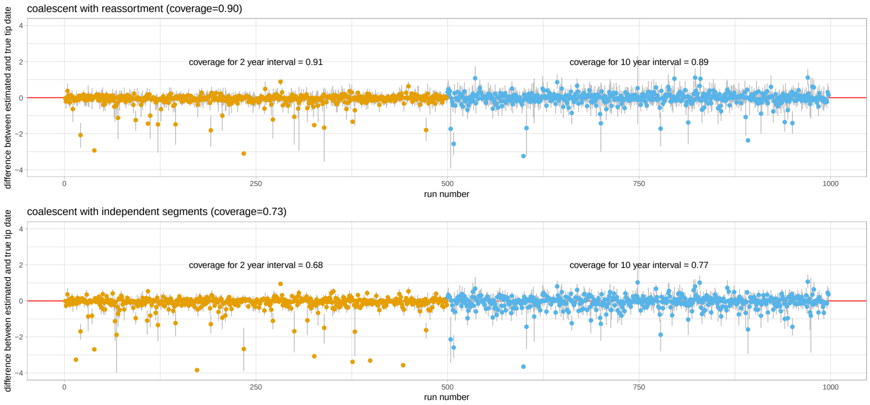


Figure S2: **Estimates of taxon sampling times of influenza A/H3N2 when using the coalescent with reassortment and the normal coalescent.** Here, we estimate the sampling times of influenza A/H3N2 taxa jointly with the phylogenetic networks respectively trees and all evolutionary parameters. To do so, we compiled 2 lots of 500 datasets with influenza A/H3N2 sequences. The first 500 datasets, were each composed of 20 genomes randomly sampled from a 2 year interval between 1995 and 2019. In the second 500 datasets, were each composed of 20 genomes randomly sampled from a 10 year interval between 1995 and 2019. For each of these datasets we randomly selected a single genome and estimated its sampling time alongside all other parameters. The figure shows the difference between the inferred and the true sampling times when using the coalescent with reassortment (top) or assuming that all segment tree are a independent realization of the sample coalescent process (bottom). The coalescent with reassortment contains the correct sampling time in the 95% highest posterior density interval in 90 % of cases. The coalescent assuming independent segment contains the true sampling time only in 68% of all cases.

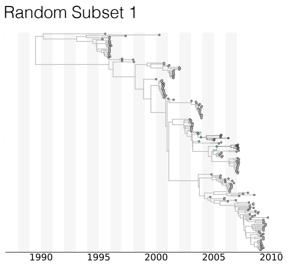


Figure S3: **Maximal clade credibility network of pandemic 1918 like influenza A/H1N1 for 3 different random subsets.**

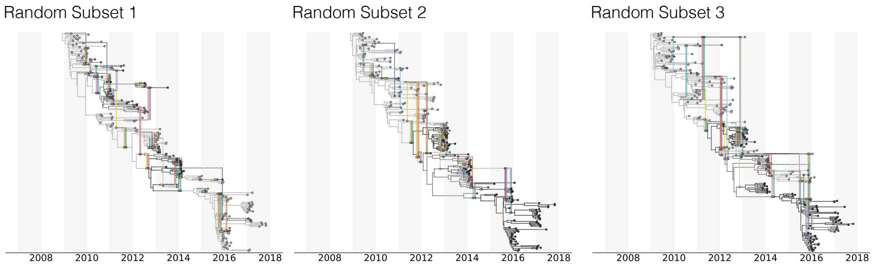


Figure S4: **Maximal clade credibility network of pandemic 2009 like influenza A/H1N1 for 3 different random subsets.**

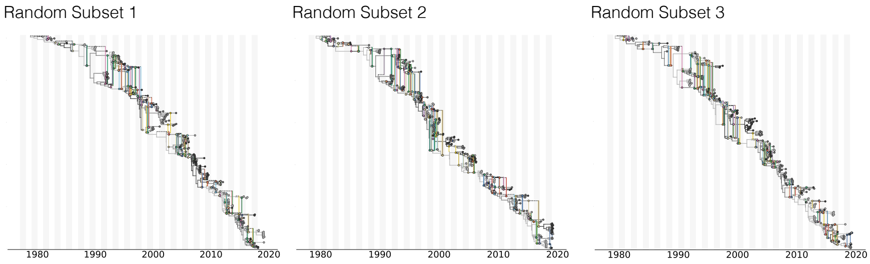


Figure S5: **Maximal clade credibility network of influenza A/H3N2 for 3 different random subsets.**

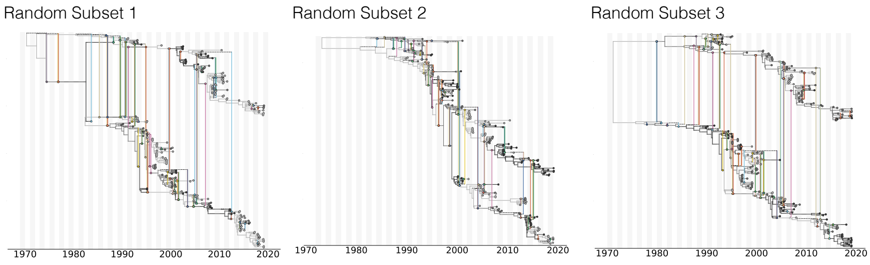


Figure S6: **Maximal clade credibility network of influenza B for 3 different random subsets.**

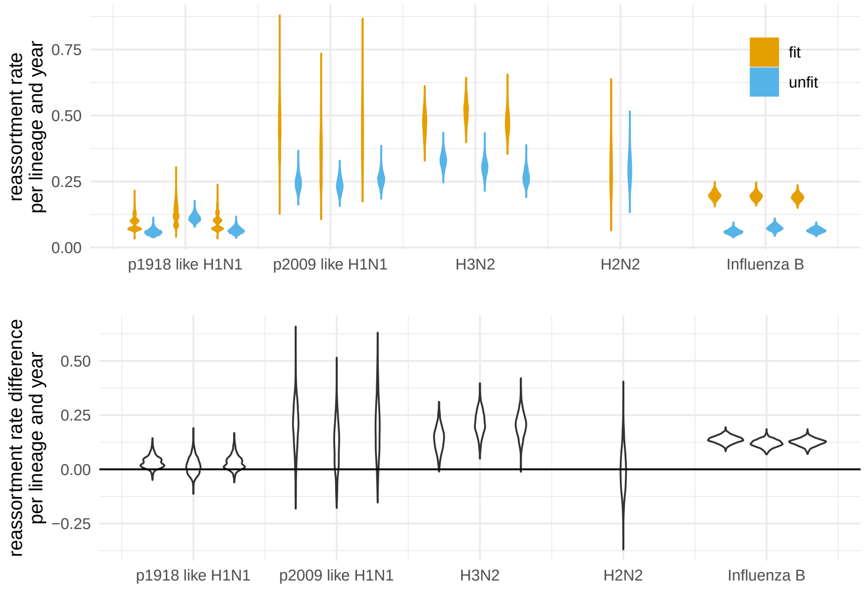


Figure S7: **Estimates of reassortment rates on fit and unfit edges when defining fit to have descendents 4 years into the future. A** Here we show the number of reassortment events on fit and unfit edges of the networks divided by the total length of fit and unfit edges. Fit edges are defined as having sampled descendent at least 4 years into the future. Every other edges is considered unfit. These rates are shown for different human influenza viruses on the x-axis. The violin plots denote the distribution of theses ratios over the posterior distribution of networks. **B** Here we show the difference between the reassortment rates on fit and unfit edges. Values above 0 indicate that reassortment events are more likely to occur on fitter, while values below 0 indicate that reassortment events are more likely to occur less fit edges.

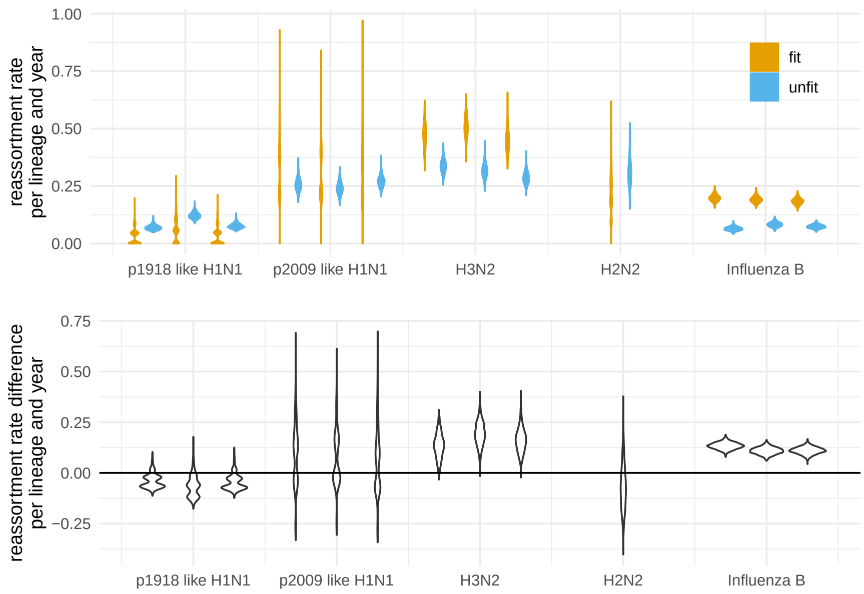


Figure S8: **Estimates of reassortment rates on fit and unfit edges when defining fit to have descendents 6 years into the future. A** Here we show the number of reassortment events on fit and unfit edges of the networks divided by the total length of fit and unfit edges. Fit edges are defined as having sampled descendent at least 6 years into the future. Every other edges is considered unfit. These rates are shown for different human influenza viruses on the x-axis. The violin plots denote the distribution of theses ratios over the posterior distribution of networks. **B** Here we show the difference between the reassortment rates on fit and unfit edges. Values above 0 indicate that reassortment events are more likely to occur on fitter, while values below 0 indicate that reassortment events are more likely to occur less fit edges.

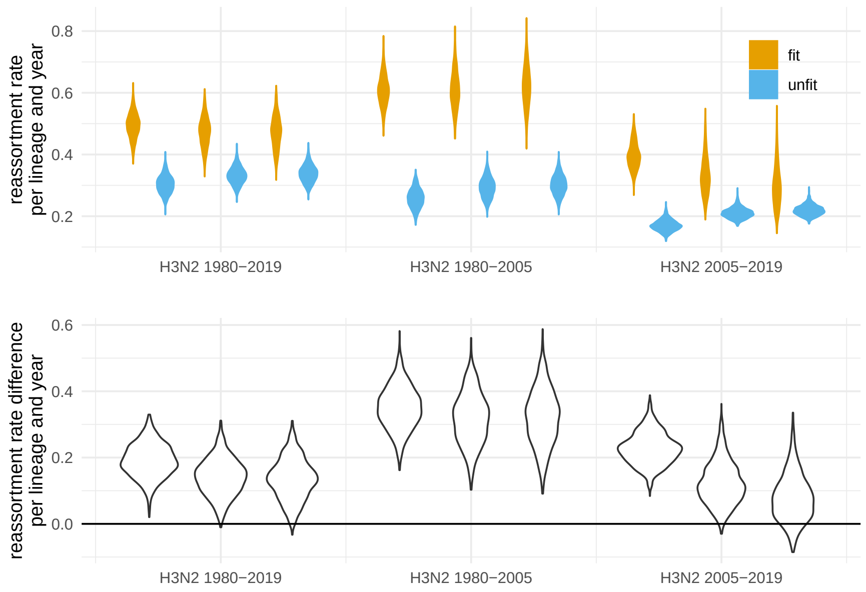


Figure S9: **Estimates of reassortment rates on fit and unfit edges for H3N2 in different time periods.** **A** Here we show the number of reassortment events on fit and unfit edges of the networks divided by the total length of fit and unfit edges for influenza A/H3N2 sampled over different timespans. The different violin plot for the same timespan show the estimates of reassortment rates or reassortment rate differences when defining a fit edges to have descendents 2,4 or 6 years into the future (from left to right) . The violin plots denote the distribution of theses ratios over the posterior distribution of networks. These rates are shown for human influenza A/H3N2 using samples at different time intervals. **B** Here we show the difference between the reassortment rates on fit and unfit edges. Values above 0 indicate that reassortment events are more likely to occur on trunk edges, while values below 0 indicate that reassortment events are more likely to occur on off-trunk edges.

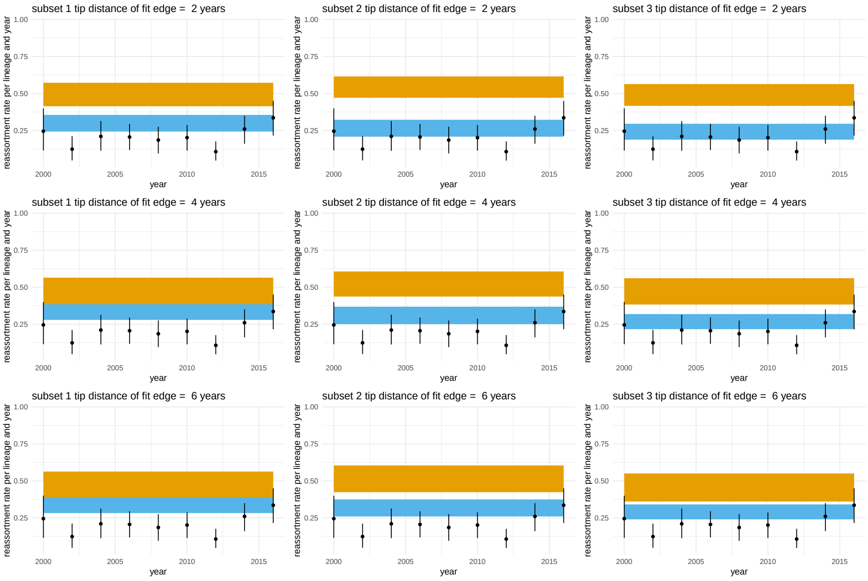


Figure S10: **Estimates of reassortment rates using influenza A/H3N2 datasets sampled over 2 years.** Here we show the estimates of reassortment rates sampled over 2 years, starting from the year on the x-axis. The orange bands show the long term estimates of reassortment rates on the fit edges for the three different random subsets (left to right) and for different definitions of what is a fit edge. From top to bottom, fit edges are defined as edges that have descendants at least 2, 4 or 6 years into the future. The blue bands show the long term estimates of reassortment rates for unfit edges.

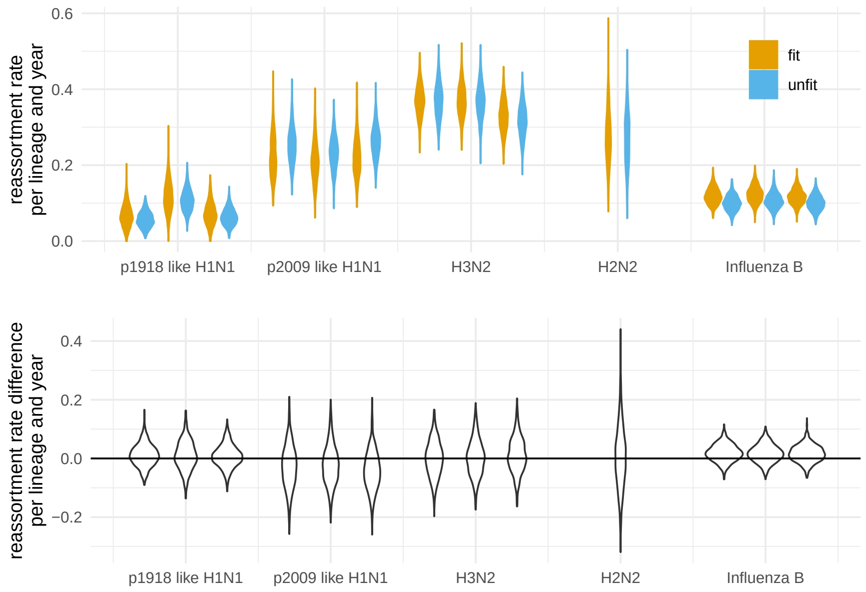


Figure S11: **Estimates of reassortment rates on fit and unfit by using posterior predictive simulations. A** Here we show the number of reassortment events on fit and unfit edges of the networks divided by the total length of fit and unfit edges computed using simulated data. The simulations use true tip sampling times of the corresponding real datasets as well as the inferred mean reassortment rates and effective population sizes to simulate networks. Fit edges are defined as having sampled descendent at least 2 years into the future. Every other edge is considered unfit. These rates are shown for different human influenza viruses on the x-axis. The violin plots denote the distribution of theses ratios over the posterior distribution of networks. **B** Here we show the difference between the reassortment rates on fit and unfit edges. Values above 0 indicate that reassortment events are more likely to occur on fitter, while values below 0 indicate that reassortment events are more likely to occur less fit edges.

### Network operators

In this section, we will describe the different network operations in more detail than in the main text.

#### Segment diversion operator

The segment diversion operator changes the path segments take at reassortment events, to do so, we follow the following procedure:

1. Choose a random reassortment edge.
2. Choose a random subset of segments for which the path at the reassortment event should be changed.
3. Remove each segment which should take another path from all ancestral network edges until a coalescent event with another edge that carries the same segment is reached.
4. Add these segments to the other parent edge of the reassortment event and follow its new path until a coalescent event with another edge that already carries the segment is reached.

#### Add/Remove operator

The Add/Remove Operator adds and removes reassortment nodes from existing networks. We first randomly choose whether to add or to remove a reassortment event. If we chose to add a reassortment event, we follow the following procedure.

1. Choose a random edge of the existing network that carries more than 1 segment.
2. Add a reassortment event on this edge with a height that is chosen uniformly at random.
3. Reattach the reassortment event on another random edge of the network and reject the move if the height of the reattachment point is lower than the height of the reassortment event.
4. Divert a random subset of the segments carried by the edge through the new network as described for the segment diversion. operator.

If we chose to remove a reassortment event, we follow the following procedure.

1. Choose a random reassortment event.
2. Choose one of the two parent edges at random to be removed.
3. Divert all segments from the edge that will be removed onto the other parent edge as described for the segment diversion.
4. Remove the parent edge and the reassortment event.

The Add/Remove operator constitutes a universal operator in that it allows us to explore all possible networks and segment trees without requiring any other operator. The following operators we describe here are more specialized operators in that they allow us to transition between certain networks more efficiently than the Add/Remove operator.

### Network scale operator

This operator scale the height of every node in the network by a random factor. Additionally, it allows to jointly scale parameters, such as the clock rate, reassortment rate or effective population size.

### Sub-network exchange

We implemented two analogues to the sub-tree exchange operators implemented in BEAST2. First, we implemented the narrow sub-network exchange operator, which is a local move. Additionally, we implemented the wide sub-network exchange operator, where move can happen over the whole network.

If we choose to perform the wide sub-network exchange, we follow the following procedure:

1. Choose two random edges of the existing network which have coalescent parent nodes.
2. Reject the move if child or parent nodes of one edge are equal to parent or child nodes of the other edge or if the height of either child node is bigger than the height of either parent node.
3. For the selected edges, remove all segments from their ancestral network edges until a coalescent event with another edge that carries the same segments is reached.
4. Detach each selected edge from its parent and reatach it at the parent node of the other edge.
5. Add all segments of each selected edge to its new parent edge. Follow the segment's path until a coalescent event with another edge that already carries the segment is reached.

If we choose to perform the narrow sub-network exchange, we follow the following procedure:

1. Choose two random edges of the exsiting network which have a common parent node.
2. Label the edge with smaller child node height $j$ and the other $p$. Label the parent edge of selected edges $jp$.
3. Reject the move if child node of the edge $p$ is not a coalescent node.
4. Choose a child edge of $p$ at random and label it $i$.
5. For edges $i$ and $j$ remove segments from the ancestral network as described for wide sub-network exchange.
6. Detach each edges $i$ and $j$ from their respective parent edges $p$ and $jp$. Reattach edge $i$ to its new parent $jp$ and edge $j$ to $p$.
7. Add segments of edges $i$ and $j$ to their ancestral network as described for widesub-network exchange.

### Sub-network slide

Next, we implemented the analogue to the sub-tree slide operation implemented in BEAST2. This operator allows to rescale the height of internal nodes while also allowing to change the topology. To do so, we follow the following procedure:

1. Choose a random edge of the exsiting network which parent is a coalescent node. Lebel this edge $i$, its parent edge $ip$, its sibling edge $j$.

2. Choose a random number $\delta$ from either uniform or Gaussian distribution. If the parent node height of edge $i$ is $t$, then its new height is $t' = t + \delta$.

3. If $\delta > 0$ , the sub-network slides upwards:

   (a) If the parent node height of edge $ip$ is larger than $t'$, set parent node height of the edge $i$ to $t'$. The network topology will not change.

   (b) Otherwise, follow the ancestal network of the edge $i$ upwards. When passing a reassortment node, randomly choose one of the two parent edges to follow. Stop when height $t'$ is reached and set it as the new parent node height of the edge $i$.

4. If $\delta < 0$ , the sub-network slides downwards:

   (a) Reject the move, if the child node height of edge $i$ is larger than $t'$.

   (b) If the child node height of the edge $j$ is smaller than $t'$, set the parent node height of the edge $i$ to $t'$. The network topology will not change.

   (c) Otherwise, detach the edge $i$ from its parent node and follow the sub-network, which is rooted at the edge $j$, downwards. When passing a coalescent node, randomly choose one of the two child edges to follow. Stop when height $t'$ is reached and set it as the new parent node height of the edge $i$.

5. Add all segments of the edge $i$ to its new parent edge. Follow the segment's path until a coalescent event with another edge that already carries the segment is reached.

6. Remove all segments of the edge $i$ from $ip$ and its ancestral network edges, until a coalescent event with another edge that carries the same segments is reached.

### Gibbs operator above the root

This operator is used for one scenario specifically, which without it can lead to massive convergence issues. Since we require networks to be rooted, part of the network can be without any information from genetic sequence data. These parts of the network are parts of the network that lie above every root of the segment trees (see figure S12). During the MCMC, these parts of the network are essentially sampled under the prior, which can be time consuming using for example Add/Remove operations. Instead of sampling these parts under the prior, we simulate under the prior instead. To do so, we do the following:

1. Check if there are any parts of the network with lie above every root of every segment. If not, do not do anything.

2. If yes, remove all edges which lie above the root of the segment tree with the largest tree height.

3. Re-simulate the sub-network starting from the edges which were remove.

Since the contribution of the Hastings-Ratio and the network probability cancel out in this move, this move will always be accepted and therefore constitutes a Gibbs move.

#### Empty segment add/remove

One of the main issues when operating on networks, is the condition that all edges in the network have to carry at least 1 segment. While this allows us to directly integrate over reassortment events that do not leave a genetic footprint in the presents, it can cause slow convergence. Operations on the network can often lead to edges carrying less than 1 segment upstream of the edge on which the actual operation was performed (see figure **??**C).

In order to enable for efficient exploration if network space, we temporarily allow networks to carry edges with 0 segments for the duration of an operation. To do so, we add a Poisson random number of empty edges to the network before any operation (fig S13A to B). We then perform an operation on a network edge that carry at least one segment (fig S13B to C). At the end we remove all empty edges in the network again (fig S13C to D) in order to only have edges with at least 1 segment left.

### Validation of MCMC operators

To validate the correctness of the coalescent with reassortment MCMC sampler, we first compare the length and height of networks and the number of reassortment events sampled from the network prior using MCMC to the statistics of networks when simulating under the same parameters. If the sampling of reassortment networks using MCMC is correct, these statistics have to be distributed the same way as when simulating networks under the same parameters.

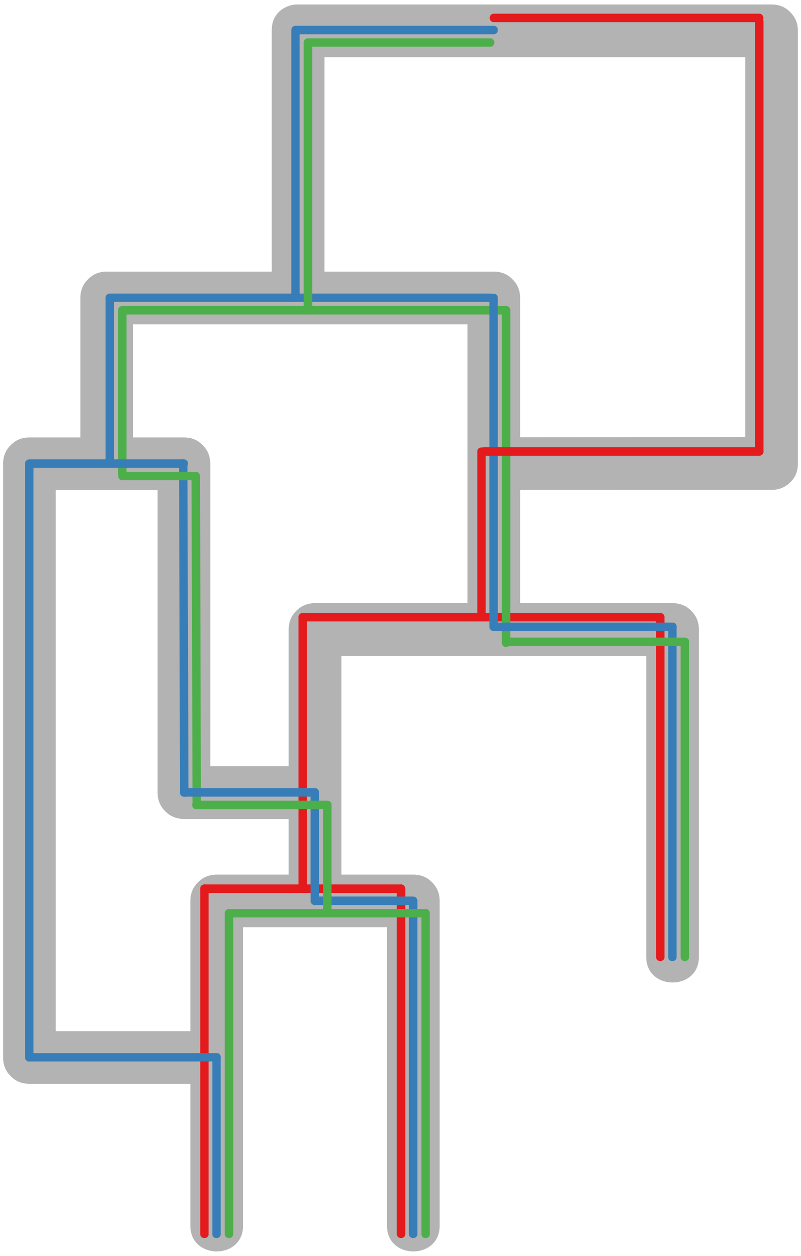

Figure S12: **Network for which parts are not informed by any genetic data.** Here, we show a network with edges which are not directly informed by genetic data. These are edges of the network which only contain segments for which the root of the segment tree are closer to the present than the network edge. The root of the network for example does contain any coalescent event in any of the three segments.

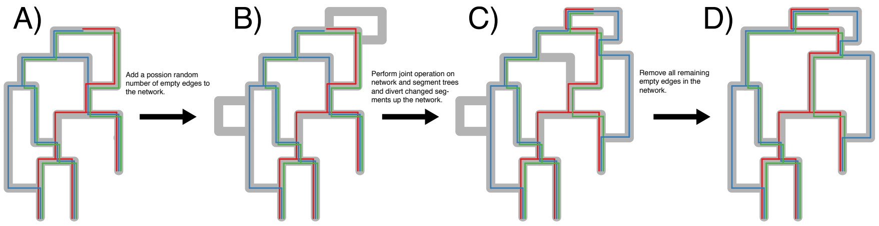


Figure S13: **Empty network edges are temporarily added to network in order to facilitate efficient exploration of network space.** Here we show how we perform joint MCMC operation on networks an the embedding of segments trees. Before we perform operations that involve segment trees, we add a poisson random number of empty network edges (A-¿B). Next, we then perform an operation on a network edge that carries segments. Here, we add a reassortment event to the network and then divert a random subset of the segments (here just the blue segment) through the new edge. We then track the blue segment up the network. At every reassortment event, we randomly choose to follow the left or the right path. We do so, until we reach a coalescent node that already carries a blue segment. In the end, we remove every empty edge in the network again.

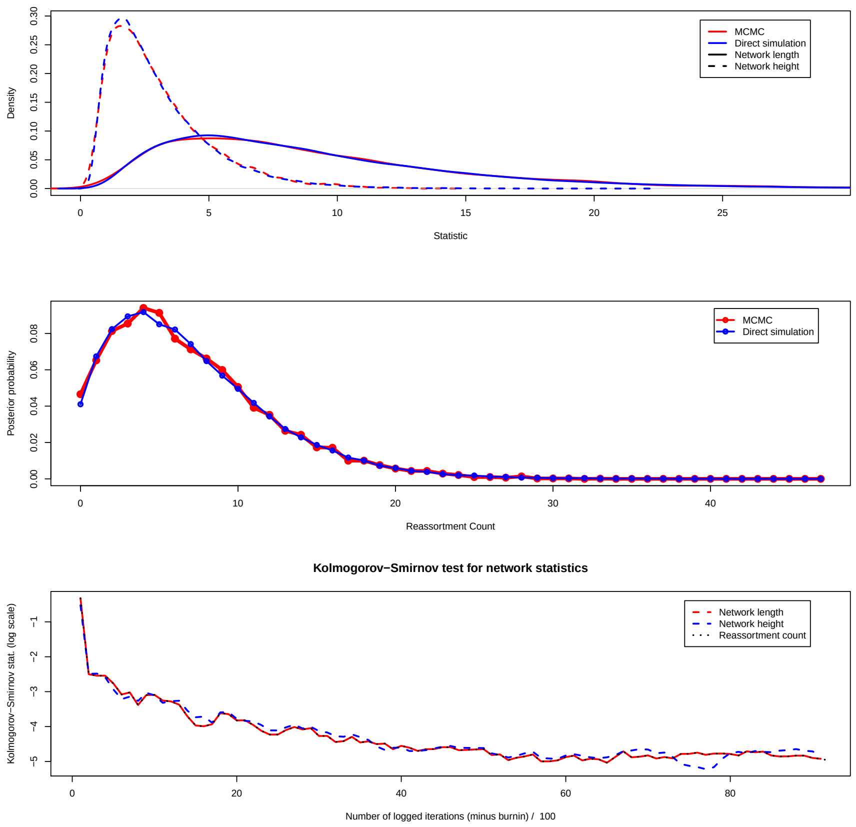


Figure S14: **Comparison between distributions inferred under the coalescent with reassortment and simulated under the coalescent with reassortment, when only using the Add/Remove operator. Top** Comparison between simulated and inferred network heights and network length. **Middle** Comparison between simulated and inferred number of reassortment events. **Bottom** KolmogorovSmirnov distance between simulated and inferred distribution over the course of the inference.

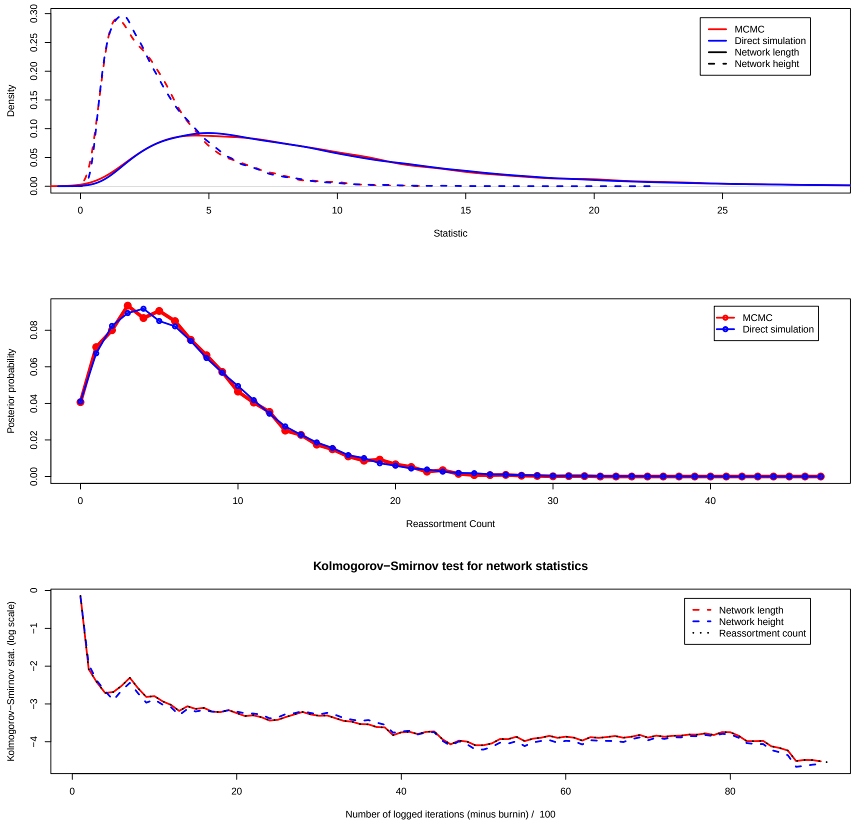


Figure S15: **Comparison between distributions inferred under the coalescent with reassortment and simulated under the coalescent with reassortment, when using the Add/Remove operator and the segment diversion operator.** In contrast to the Add/Remove operator, the segment diversion operator is not a universal operator and in order to explore any possible network, we run the comparison using both operators **Top** Comparison between simulated and inferred network heights and network length. **Middle** Comparison between simulated and inferred number of reassortment events. **Bottom** KolmogorovSmirnov distance between simulated and inferred distribution over the course of the inference.

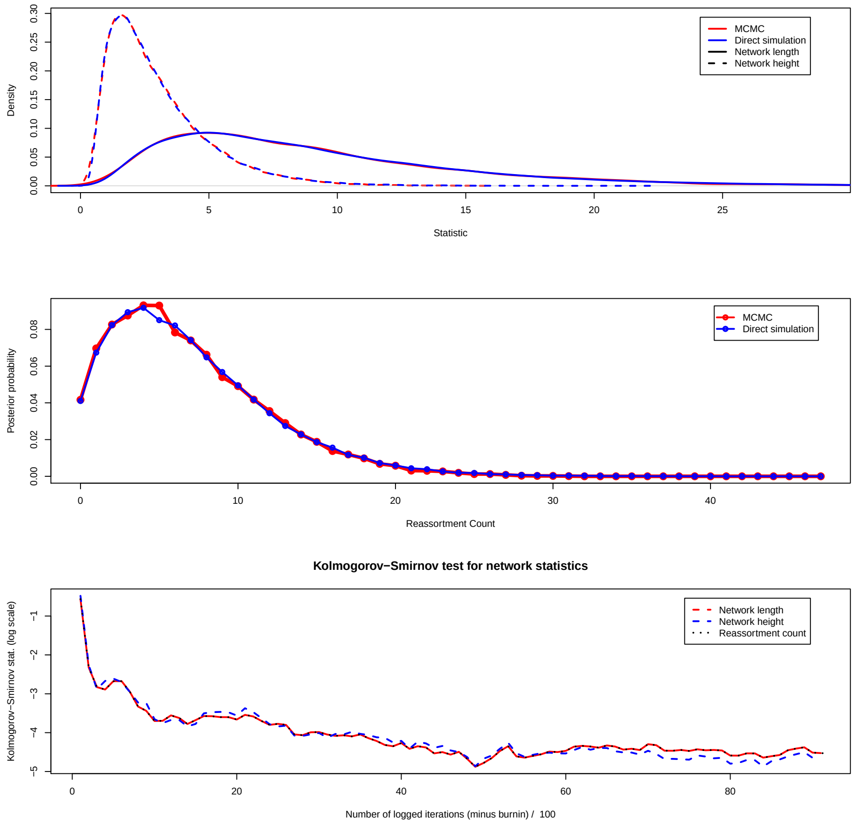


Figure S16: **Comparison between distributions inferred under the coalescent with reassortment and simulated under the coalescent with reassortment, when using the Add/Remove operator and the network scale operator.** In contrast to the Add/Remove operator, the network scale operator is not a universal operator and in order to explore any possible network, we run the comparison using both operators **Top** Comparison between simulated and inferred network heights and network length. **Middle** Comparison between simulated and inferred number of reassortment events. **Bottom** KolmogorovSmirnov distance between simulated and inferred distribution over the course of the inference.

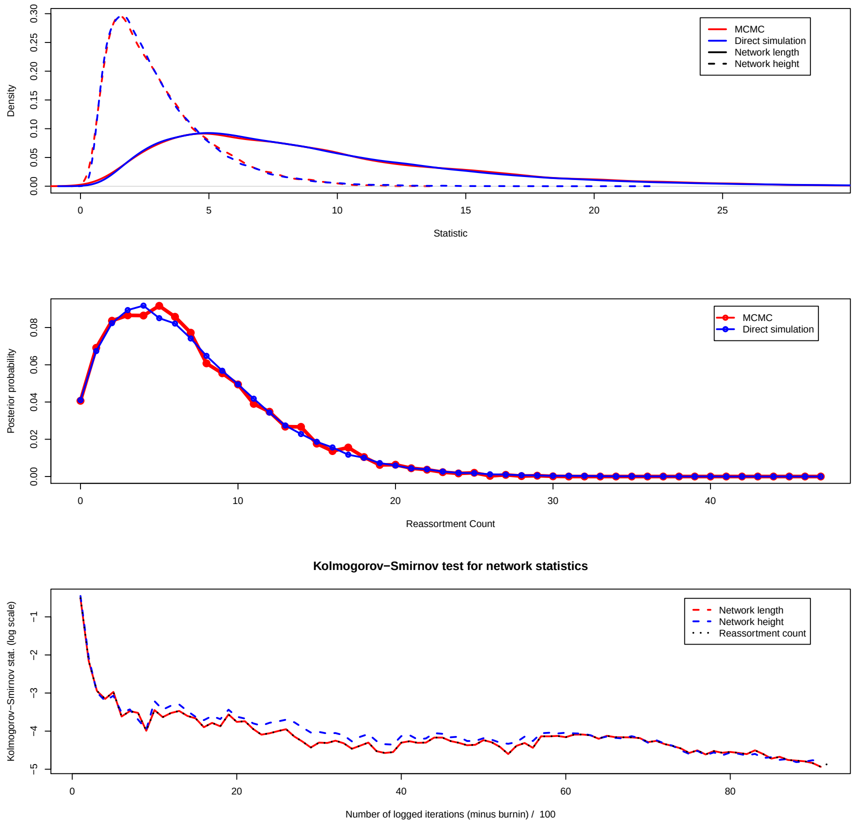


Figure S17: **Comparison between distributions inferred under the coalescent with reassortment and simulated under the coalescent with reassortment, when using the Add/Remove operator and the wide sub-network exchange operator.** In contrast to the Add/Remove operator, the wide sub-network operator is not a universal operator and in order to explore any possible network, we run the comparison using both operators **Top** Comparison between simulated and inferred network heights and network length. **Middle** Comparison between simulated and inferred number of reassortment events. **Bottom** KolmogorovSmirnov distance between simulated and inferred distribution over the course of the inference.

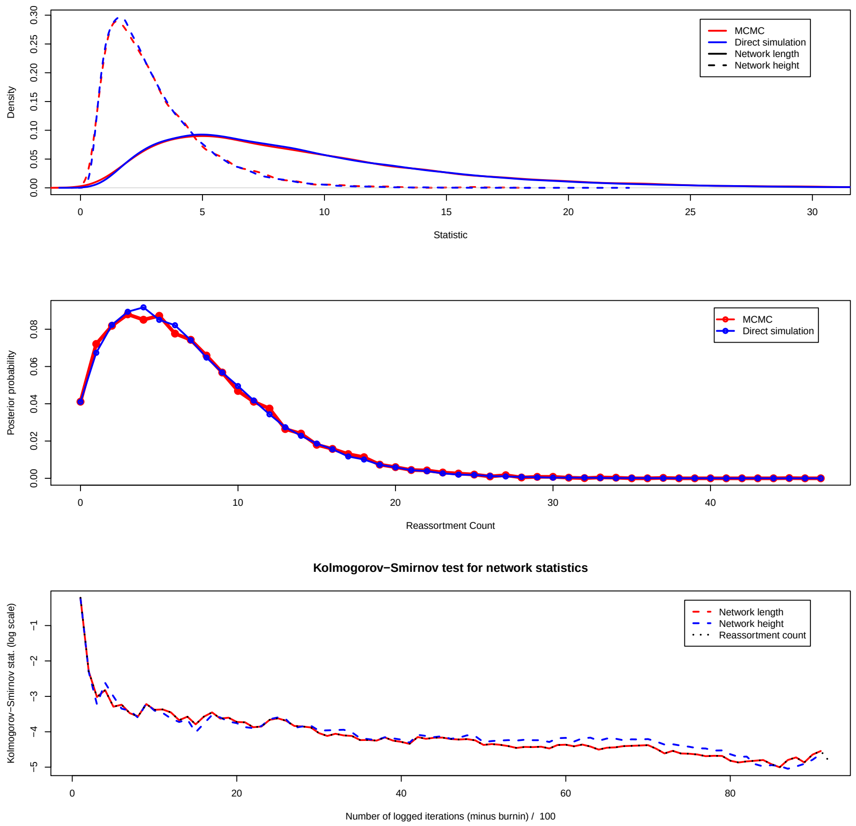


Figure S18: **Comparison between distributions inferred under the coalescent with reassortment and simulated under the coalescent with reassortment, when using the Add/Remove operator and the narrow sub-network exchange operator.** In contrast to the Add/Remove operator, the narrow sub-network operator is not a universal operator and in order to explore any possible network, we run the comparison using both operators **Top** Comparison between simulated and inferred network heights and network length. **Middle** Comparison between simulated and inferred number of reassortment events. **Bottom** KolmogorovSmirnov distance between simulated and inferred distribution over the course of the inference.
